# Neuro-immune Crosstalk in the Enteric Nervous System from Early Postnatal Development to Adulthood

**DOI:** 10.1101/2022.05.12.491517

**Authors:** Viola Maria Francesca, Chavero-Pieres Marta, Modave Elodie, Stakenborg Nathalie, Delfini Marcello, Naomi Fabre, Iris Appeltans, Tobie Martens, Katy Vandereyken, Jens Van Herck, Philippe Petry, Simon Verheijden, Sebastiaan De Schepper, Alejandro Sifrim, Katrin Kierdorf, Marco Prinz, Pieter Vanden Berghe, Thierry Voet, Guy Boeckxstaens

## Abstract

Correct development and maturation of the enteric nervous system (ENS) is critical for survival. Early in life, the ENS requires significant refinement in order to adapt to the evolving needs of the tissue, changing from milk to solid food at the time of weaning. Here, we demonstrate that resident macrophages of the muscularis externa, MMϕ, refine the ENS early in life by pruning synapses and phagocytosing abundant enteric neurons. After weaning, MMϕ continue to closely interact with the ENS, acquire a microglia-like phenotype and are crucial for the survival of enteric neurons. Of note, this microglia-like phenotype is instructed by TGFβ produced by the ENS, introducing a novel reciprocal cell-cell communication responsible for the maintenance of the neuron-associated MMФ niche in the gut. These findings elucidate a novel role of intestinal macrophages in ENS refinement early in life, and open new opportunities to treat intestinal neurodegenerative disorders by manipulating the ENS-macrophage niche.

## Introduction

In mammals, gut function is largely controlled by the enteric nervous system (ENS), a complex and vast collection of neurons organized in two plexuses, the myenteric and submucous plexus. Even in the absence of input from the central nervous system (CNS), the ENS efficiently coordinates a plethora of vital functions including nutrient absorption, peristalsis and intestinal secretion^1^. It has been estimated that the adult human ENS contains over half a billion neurons, and therefore is often referred to as the “brain of the gut”. Diseases of the ENS range from limiting life-quality to lethal, and can arise during early development or manifest as neurodegeneration later in life^2–4^.

The ENS develops from progenitors of the neural crest, which colonise the gastrointestinal tract prior to birth in a process that is extremely fine-tuned in time and space, and is orchestrated by an interplay between signalling molecules and receptors that ensures complete innervation of the gut^5^. While the process of colonisation by progenitors terminates prior to birth, neuronal subtypes develop at differing rates, and the full range of neuronal differentiation is only reached after birth^6^. In the ENS, neuronal maturation persists after birth, with changes in neurochemical composition, soma size and synaptic input reported to occur between early postnatal life and adulthood^7^. In line, the randomly occurring contractions required for motility only appear several days after birth, and reach a mature frequency and amplitude after weaning^8^. This maturation pattern appears to mirror the developing need of the intestine to first propel liquid, and only subsequently solids, through the gastrointestinal tract. Indeed in mouse, experiments show that the intestine is able only after postnatal day (P) 14 to expel beads, and the speed at which this occurs increases significantly from P14 and P21^8^. These findings suggest that the ENS undergoes significant maturation in early postnatal life, however the mechanisms involved in this process have yet to be elucidated.

Tissue-resident macrophages are highly specialized phagocytes that actively contribute to immune defence but also to tissue homeostasis of their host organ^9,10^.This task relies on their ability to sense and respond to local demands including metabolic changes, tissue damage and microbial insults, while performing tissue-specific functions to support surrounding cells. The instruction of the tissue resident macrophage comes largely from the surrounding cells, the ‘macrophage niche’, that provide the molecular cues to initiate a niche-specific transcriptional profile^11^. Thus, tissue-resident macrophages represent highly specialised cells that adapt functionally and transcriptionally to the tissue in which they reside. Furthermore, macrophages display a certain degree of plasticity, enabling transition of phenotype in response to altered environmental needs, in particular in the context of inflammation^12^. The perhaps most striking example of macrophage plasticity is the tissue-resident macrophage population in the brain parenchyma, microglia. Microglia serve a plethora of specific functions involved in neuronal formation, maturation and wiring during defined developmental time windows, such as synaptic pruning and the shaping of neuronal circuits during postnatal development or trophic support and regulation of adult hippocampal neurogenesis in the adult^13–15^.

In the gut, tissue-resident macrophages are present throughout its different layers, and carry out distinct and specialised functions according to their specific anatomical location^16^. We recently identified a subpopulation of long-lived resident macrophages in the muscularis externa that lies in close association with enteric neurons in the myenteric plexus, whose health and survival are critical for gastrointestinal motility^17^. These muscularis macrophages (MMФ) receive a variety of signals from the surrounding neurons, ranging from colony stimulating factor 1 (Csf1) which is essential for their survival, to adrenergic input installing a tolerogenic phenotype in the context of inflammation and infection^18,19^. Conversely, depletion of this MMФ population in adulthood leads to neurodegeneration, delayed intestinal transit, impaired intestinal motility and colonic dilation^17^. While the role of MMФ in neuronal homeostasis in adulthood has been clarified, it is unclear what role MMФ play during early postnatal development. Here, we show that MMФ, similar to microglia in the CNS parenchyma, contribute to the maturation of the ENS early in life. Furthermore, we identified the time of weaning as crucial ‘tipping-point’ for macrophage function, at which MMФ shift their role from neuronal refinement to neuronal support. This shift is accompanied by profound transcriptional and phenotypic maturation imprinted by transforming growth factor β (TGFβ) produced by the ENS, introducing a novel cell-cell circuit responsible for the maintenance of the neuron-associated MMФ niche in the gut. Taken together, these findings elucidate a novel role of intestinal macrophages in ENS refinement early in life, and open new opportunities to modulate and target the diseased ENS-macrophage niche.

## Results

### MMФ refine the ENS during early postnatal development

Previous reports have suggested that the ENS undergoes remodelling during early postnatal development, including changes of soma size, organization of ganglia and synaptic density^7,8^. To carefully define these age-related changes in ENS architecture, we first assessed neuronal density during early postnatal development (P10), at weaning (P21) and during adulthood (P56). At P10, neuronal cell bodies were abundant throughout the plane between the circular and longitudinal muscle layers, loosely organised into ganglia that run parallel to the circular muscle layer, while delimitations between ganglia were ill-defined and interconnecting fibres were disorganised (Figure 1A). Density of enteric neurons, ganglia and single extra-ganglionic neurons dynamically decreased from postnatal development to adulthood, also when tissue growth was taken into account (Figure 1A-D, S1A-C). These changes were accompanied by a decrease in interconnecting fibres (Figure 1E). Density of the presynaptic marker Synapsin I was measured at P7, P14, P21 and P56, with a significant increase at P14 compared to younger (P7) and older (P21, P56) timepoints (Figure 1F)^7^. These findings suggest that the early postnatal phase prior to weaning represents a dynamic phase for the remodelling and the refinement of the enteric neuronal network.

**Figure 1:**
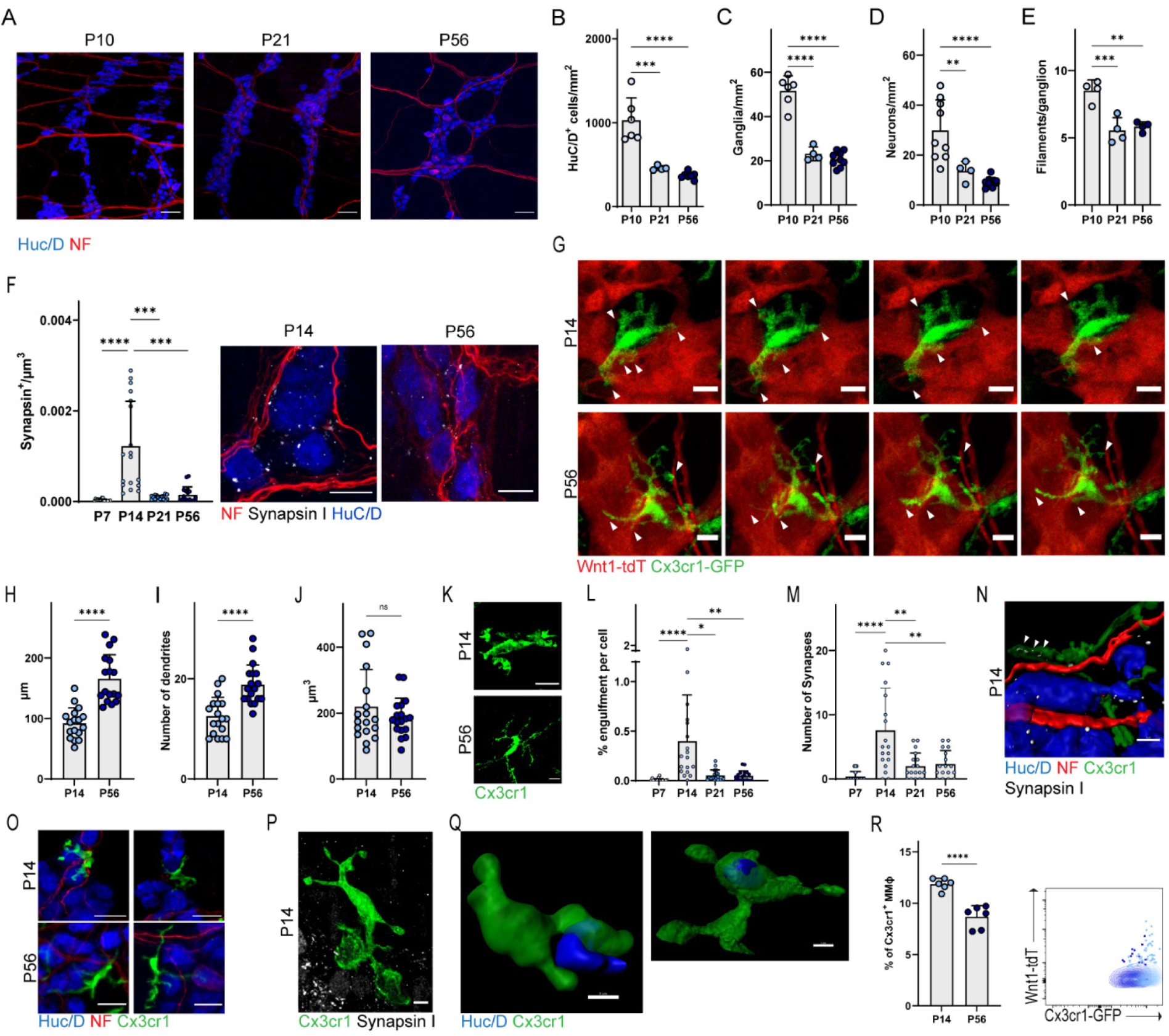
MMϕ prune synapses and engulf neurons during early postnatal development. (**A**) Representative confocal images of the myenteric plexus at P10, P21 and P56. Scale bar is 50µm. Quantification of neuronal density (**B**), density of ganglia (**C**), single extra-ganglionic neurons (D) and interconnecting fibre bundles (**E**) in the myenteric plexus at P10, P21 and P56. (**F**) Quantification of the volume of Synapsin1 per µm3. Representative confocal image of myenteric neurons at P14 and P56. Scale bar is 10µm. (**G**) Time-lapse stills taken from ex vivo live-imaging of the muscularis externa of 14 and 56 day-old Wnt1^Cre/WT^ Rosa26^tdT/WT^ Cx3cr1^GFP/WT^ mice. Arrows indicate phagocytic cups and filopodia of MMϕ extending and retracting towards neural cells. Scale bar is 10µm. Total length of MMϕ filopodia (**H**), number of dendrites (**I**) and MMϕ cell volume (**J**) as assessed via 3D reconstruction using Imaris. (**K**) Representative confocal images displaying MMϕ morphology at P14 and at P56. Scale bar is 10µm. (**L**) Percentage of synaptic engulfment per cell, determined as volume of Synapsin^+^ volume/MMϕ volume. (**M**) Number of Synapsin I+ puncta engulfed by MMϕ, determined via digital reconstruction with Imaris. (**N**) Digital reconstruction using Imaris depicting engulfment of synapses by MMϕ. White arrowheads indicate engulfed synapses. (**O**) Representative confocal images displaying MMϕ engulfing neuronal cell bodies and extending filopodia towards neuronal cell bodies and filaments. Scale bar is 20µm. (**P**) Representative confocal image showing a MMϕ extend phagocytic cups toward enteric neurons. Scale bar is 2µm. (**Q**) Digital reconstructions of MMϕ engulfing enteric neurons, performed using Imaris. Scale bar is 5µm. (**R**) Quantification via flow cytometry of percentage of MMϕ engulfing neural cells (Wnt1-tdTomato^+^), and representative dotplot of Wnt1-tdTomato signal in MMϕ. Data are shown as Mean ± SD, analysed using t-test, ANOVA (followed by Tukey’s multiple comparisons) or Kruskal-Wallis (followed by Dunn’s multiple comparisons). Data are pooled from at least 2 independent experiment, apart from experiments shown in F, H-J and L-M. * *p* < 0.05; ** *p* < 0.01; *** *p* < 0.001; **** *p* < 0.0001; ns = not significant.

We hypothesized that MMϕ, like microglia in the CNS, refine the ENS during development and closely interact with enteric neurons. To assess neuron-MMϕ interaction, we generated mice expressing tdTomato in enteric neurons and green fluorescent protein (GFP) in Cx3cr1^+^ macrophages, including MMϕ, (*Cx3cr1*^*GFP/WT*^ *Wnt1*^*CRE/WT*^ *Rosa26*^*tdT/WT*^). Using ex-vivo live-imaging of the muscularis externa at P14, we observed that Cx3cr1^high^ MMϕ extend and retract phagocytic cups towards neurons (Figure 1G + supplementary videos 1 and 2). At P56, MMϕ the processes appeared more ramified and elongated towards adjacent enteric neurons, compared to the processes of MMϕ at P14, where phagosomes and vesicles were evident (figure 1G). In line, analysis of MMϕ morphology revealed a significant increase in filopodia length and number of dendrites at P56 compared to P14, while no changes in cell body volume were observed (Figure 1H-K).

As we showed that MMϕ indeed dynamically interact with enteric neurons, we next investigated whether MMϕ prune synapses in the ENS. Synaptic engulfment was assessed using Synapsin-I immunostaining and confocal imaging, followed by digital reconstruction as previously described^20^. At P14 MMϕ engulfed significantly more Synapsin I^+^ puncta compared to P21 and P56 (figure 1L-N). Furthermore, MMϕ also engulfed neuronal bodies at P14, but not anymore at P56 (Figure 1O-Q). To further assess whether MMϕ may also be engulfing neurons or ENS debris, in addition to synapses, we made use of *Cx3cr1*^*GFP/WT*^ *Wnt1*^*CRE/WT*^ *Rosa26*^*tdT/WT*^ mice, and determined neural tdTomato within Cx3cr1^+^ MMϕ via flow cytometry. The percentage of Cx3cr1^GFP^ positive MMϕ containing ENS-derived tdTomato^+^ was significantly higher at P14 compared to P56 (figure 1R, S1D). Taken together, these findings suggest that early postnatal development is a period of dynamic neuronal refinement in the ENS, and MMϕ actively participate in the remodelling of the ENS via contacting adjacent neurons and phagocytosing of neuronal bodies and synapses.

### MMФ function differs according to the developmental stage of the gut

In adulthood, we previously demonstrated that enteric neurons rely on MMϕ for their survival, as indicated by the observation that depletion of neuron-associated MMϕ leads to neurodegeneration^17^. Whereas we demonstrated above that MMϕ refine the ENS early in life, rather than already providing trophic support, we hypothesized that MMϕ depletion during the early postnatal phase might have consequences for the development and the maturation of the ENS. To test this hypothesis, mice received 2 injections of a depleting αCSF1r 3 days apart, either during postnatal development (P10), at weaning (P21), or in adulthood (P56), and were analysed 7 days after the first injection (Figure 2A). Treatment with αCSF1r led to a significant reduction in MMϕ, and did not alter other immune cell populations in the muscularis externa, including B cells and T cells (Figure S2A,B). In line with our previous findings, depletion of MMФ during adulthood led to a significant reduction in neuronal density compared to controls^17^ (Figure 2B,C). In contrast, if MMϕ were depleted during postnatal development prior to weaning (P10), neuronal density was found to be significantly increased in treated mice compared to controls (Figure 2B,C). Strikingly, ganglia maintained the morphological appearance characteristic of P10, with ill-defined ganglia and reduced distance between them, while in control mice, maturation of ganglia had continued and appeared more organized resembling ganglia at P21 (Figure 2B, C). Depletion of MMФ at weaning (P21) led to no significant changes in neuronal density or morphology of ganglia (Figure 2B, C). Taken together, our findings support a role in MMϕ in neuronal refinement during early postnatal life. Furthermore, we confirmed again the neuro-supportive role of MMϕ in adulthood, suggesting that the neuron-MMϕ interaction within the niche differs significantly at distinct stages in development.

**Figure 2:**
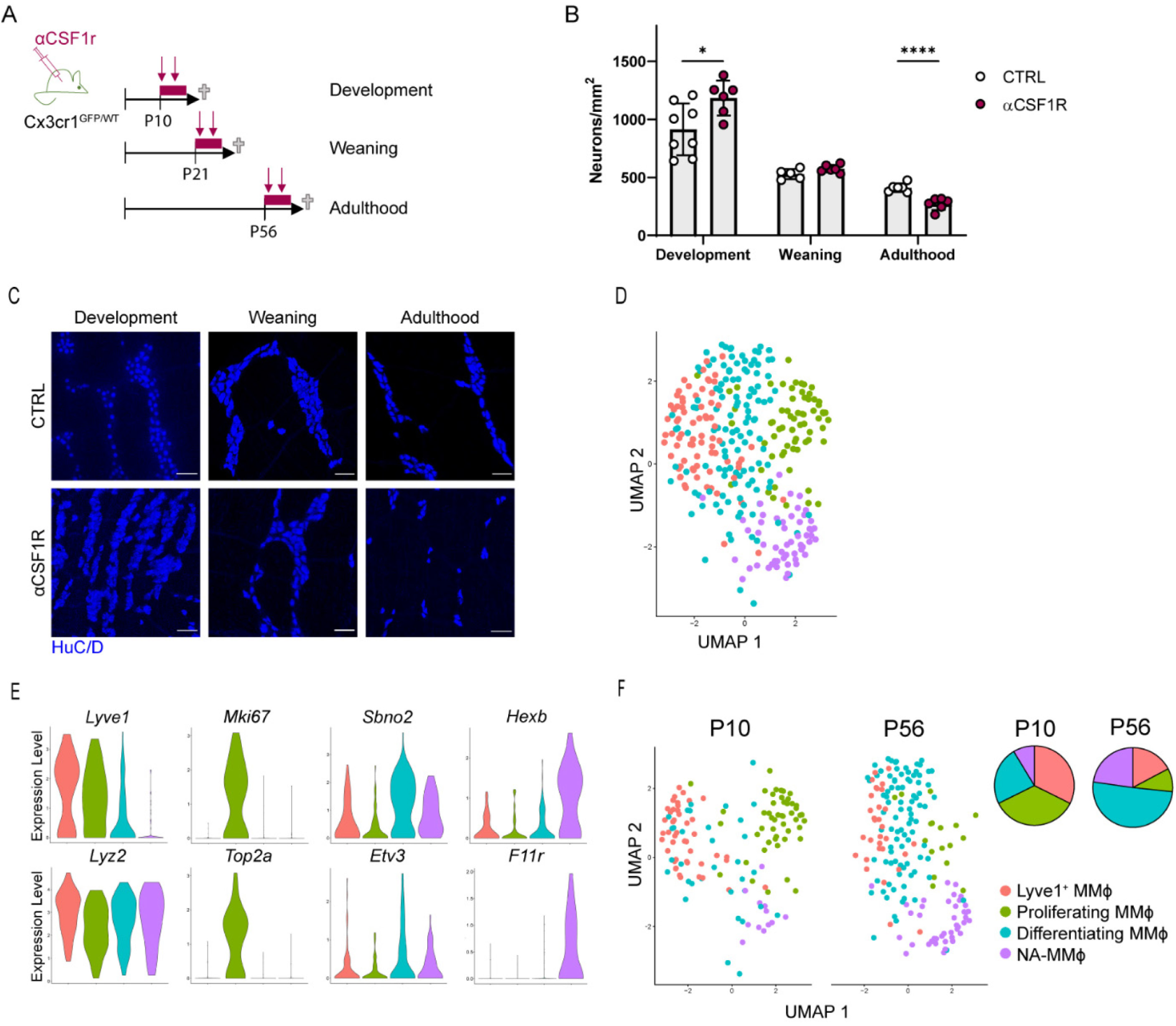
MMϕ undergo functional and transcriptional shifts from early postnatal development to adulthood. (**A**) Diagram to illustrate the treatment of mice with two injections αCSF1R, to deplete MMϕ for the duration of 7 days, during either postnatal development, weaning or adulthood. (**B**) Neuronal density in αCSF1R-treated mice and controls. Pooled data from 2 experiments. Multiple t-tests. Data are shown as Mean ± SD. Data are pooled from 2 independent experiments. (**C**) Representative confocal images of the myenteric plexus in αCSF1R-treated mice and controls. Scale bars are 50µm. (**D**) UMAP dimensionality reduction and clustering analysis of SMARTSeq2 scRNAseq data (n=319 cells) reveals four subpopulations of MMϕ, colour coded on the UMAP plot. (**E**) Violin plots displaying expression distributions of individual genes in cells grouped by cluster identified in D. (**F**) UMAPs by age, with colour coding reflecting the clusters identified in the integrated dataset.

### Gene expression profiling revealed transcriptomic adaptation of MMϕ

To investigate potential differences in the transcriptional signature underlying the functional transition of MMϕ prior to and after weaning, Cx3cr1^+^ MMϕ were FACS sorted from Cx3cr1^GFP/WT^ mice at P10 and P56 and further used for single-cell RNA sequencing using the smartSEQ2 platform (Figure S2C). We intended to use this sequencing protocol rather than the commonly used 10x Genomics Chromium platform in order to acquire high transcriptomic coverage^21^. Following quality control, we obtained a total of 353 cells, of which 169 were from P10 mice and 184 from P56 mice (Figure S2E). Unbiased hierarchical clustering revealed 5 transcriptionally distinct cellular subpopulations, which were all classified as macrophages based on their expression of signature genes including *Ptprc, Csf1r* and *Adgre1* (F4/80) (Figure S2D,E). Expression of monocyte markers such as *Ccr2* or *Ly6c2* was low or not detected (Figure S2E). One cluster, comprising 34 cells and present almost exclusively in the P10 samples, contained a high amount of transcripts of extracellular matrix genes, including *Col3a1* and *Dcn*, suggesting potential contamination of their transcriptional signatures due to phagocytosis of cells depositing the extracellular matrix (Figure S2F). At P10, some MMФ were indeed embedded within the extracellular matrix, which expressed the identified markers such as *Dcn* (Figure S2G). As these findings suggest that these cells clustered separately due to the contamination by transcripts of phagocytosed cells, we excluded this cluster from further analysis (Figure 2D).

We next annotated the clusters using the integrated dataset. The most abundant cluster was characterised by an upregulation of *Lyve1* and genes implicated in phagocytosis, including *Lyz2, Apoe* and *Ctsd*, and we thus named this cluster ‘Lyve1^+^ MMϕ’ (Figure 2E). The second cluster consisted of proliferating MMФ, and was characterised by the upregulation of cell cycle genes, including *Mki67, Top2a, Cdkn1a, Smc4* and *Tubb* (Figure 2E). The third cluster was composed of MMФ undergoing differentiation, as it was characterised by the upregulation of several transcription factors involved in terminal macrophage differentiation including *Etv3, Socs3* and *Irf1*, and transcriptional co-regulators such as *Sbno2* (Figure 2E). The fourth cluster was characterised by the expression of genes considered to be microglia signature genes, including *Hexb, Ctss, Tmem119, F11r, Olfml3, Sema4d, Cx3cr1* (Figure 2E). Upregulation of microglia signature genes has been described in macrophages associated to peripheral nerves, so we named this cluster neuron-associated MMϕ (NA-MMϕ)^22,23^.

When the abundance of each identified cluster was assessed by age, we found that the P10 sample comprised of mainly Lyve1+ MMϕ and proliferating MMϕ (Figure 2F). In the adult (P56) sample, cells were either undergoing terminal differentiation, or had acquired a NA-MMϕ transcriptional signature (Figure 2F). These findings suggest that the change in function of MMϕ is based on a profound shift in transcriptomic signature of MMϕ from postnatal development to adulthood.

### Distinct MMϕ subsets are located in different subtissular niches

Next, we aimed to further characterize the identified subclusters in terms of their anatomical localisation functions. First, we quantified the Lyve1^+^ MMϕ cluster via flow cytometry in MMϕ isolated at P10 and P56. We observed significantly more Lyve1^+^ MMϕ at P10 than at P56 (Figure 3A). As recently described, a Lyve1^hi^MHCII^lo^ macrophage subtype resides within numerous organs, including the intestine, and is associated to vasculature^24^. The Lyve1^+^ cluster identified here resembles the Lyve1^hi^MHCII^lo^ cluster transcriptionally, with upregulation of *Fcna, Lyve1, Cbr2, Cd209f, Ninj1* and *Hmox1* (Figure S3A). Similarly, these cells could be found either associated to VE-Cadherin^+^ blood vessels within the myenteric plexus, but the majority of these cells were closely associated to capillaries located within the serosa (Figure 3B). A similar anatomical distribution was observed at both P10 and P56, however Lyve1^+^ MMϕ were rather scarce at P56. As this cluster upregulated genes involved in phagocytosis, including *Apoe* and *Lyz2*, hence we hypothesised that Lyve1^+^ MMϕ may be specialised in phagocytosis. Phagocytic capacity of MMϕ was assessed via an ex-vivo phagocytosis assay using fluorescently labelled dextran. Lyve1^+^ MMϕ had a significantly higher phagocytic index than the other identified MMϕ subtypes at P56, suggesting that this cluster represents MMϕ specialised in phagocytosis and engulfment (Figure 3C, S3C).

**Figure 3:**
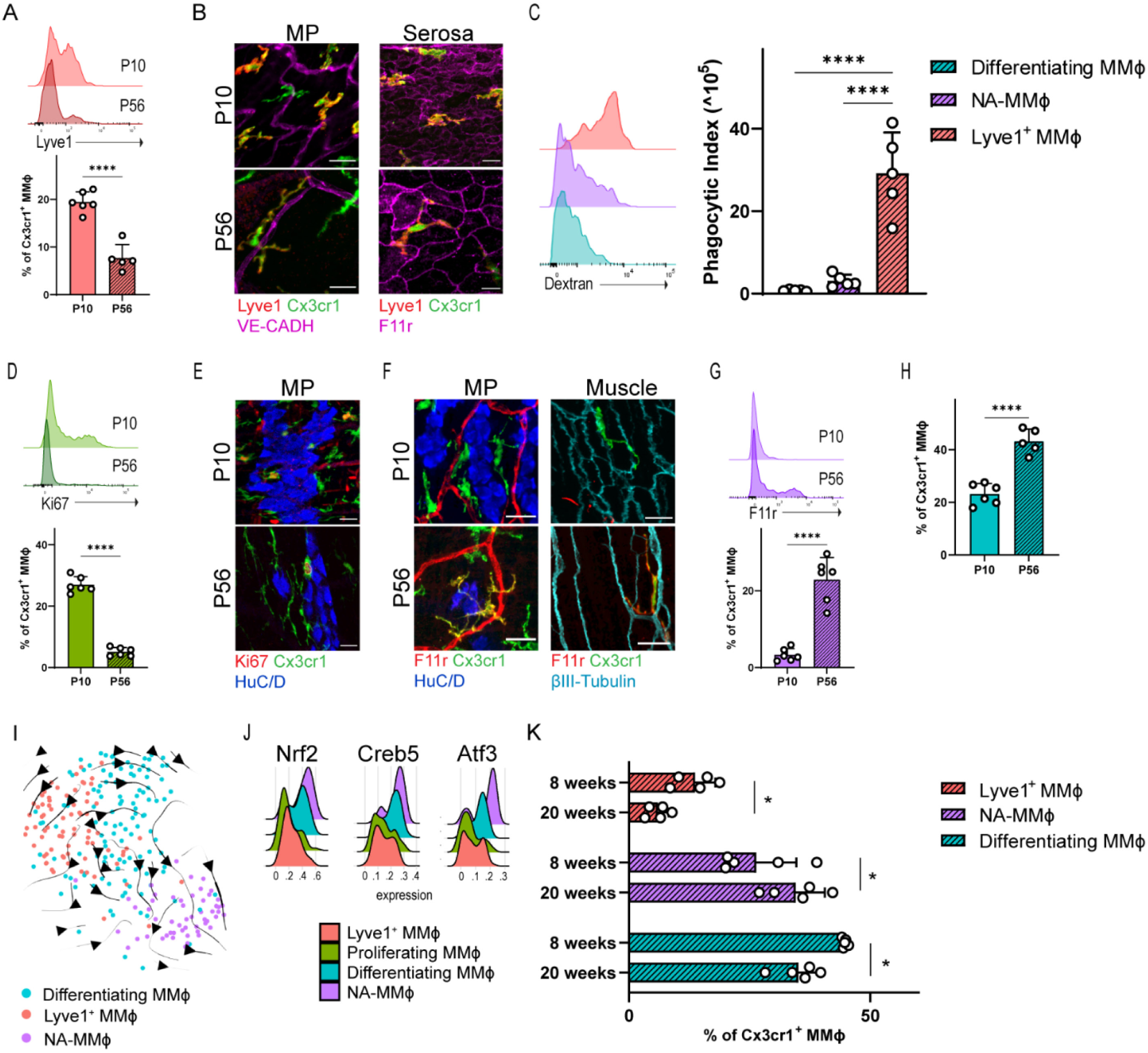
MMФ subsets occupy distinct micro-anatomical niches in the muscularis externa. (**A**) Representative histogram depicting Lyve1 expression in total MMФ, and quantification of Lyve1^+^ MMФ in the muscularis externa of P10 and P56 mice via flow cytometry. (**B**) Representative confocal images showing Lyve1^+^ MMФ in the myenteric plexus (MP) and serosa at P10 and at P56. (**C**) Quantification of phagocytic index of identified MMФ populations and representative histograms depicting fluorescent dextran signal within each population. (**D**) Representative histogram depicting Ki67 expression in total MMФ, and quantification of Ki67^+^ MMФ in the muscularis externa of P10 and P56 mice via flow cytometry. (**E**) Representative confocal images showing Ki67^+^ MMФ in the myenteric plexus (MP) at P10 and at P56. (**F**) Representative confocal images showing F11r^+^ MMФ in the myenteric plexus (MP) and the muscle layer at P10 and at P56. (**G**) Representative histogram depicting F11r expression in total MMФ, and quantification of F11r^+^ MMФ in the muscularis externa of P10 and P56 mice via flow cytometry. (**H**) Quantification via flow cytometry of Lyve1^-^F11r^-^Ki67^-^ MMФ in the muscularis externa at P10 and at P56. (**I**) RNA Velocity analysis (n=253) reveals direction of transcriptional shifts between MMФ. Proliferating MMФ were excluded from this analysis. (**J**) Ridge plots SCENIC AUC distribtuions representing the activation of transcription factors significantly upregulated in NA-MMФ (**K**) Quantification via flow cytometry of population frequency at 8 weeks (P56) and 20 weeks. In all data shown, MMФ were obtained by gating on live CD45^+^Cd11b^+^Cd64^+^Cx3cr1^+^. Data are shown as Mean ± SD and are representative of at least 2 independent experiments. Data are analysed using T-tests or ANOVA followed by Tukey’s multiple comparisons. Scale bars are 20µm. * *p* < 0.05; **** *p* < 0.0001.

The second cluster most abundant cluster at P10 was composed of proliferating MMФ. In line with the scRNAseq findings, proliferation of MMФ, assessed by intracellular Ki67 expression, was increased at P10 compared to P56 (Figure S2B, 3D). While around 30% of MMФ were undergoing replication at P10, at P56, upon completion of tissue growth, only 5% of MMФ were proliferating. Also via immunohistochemistry, Ki67^+^ MMФ could be observed in the myenteric plexus at both P10 and P56 (Figure 3E). The high degree of proliferation coupled with the absence of monocytes in the tissue suggests that the expanding muscularis niche is filled via proliferation of self-maintaining macrophages rather than by infiltration of monocytes that differentiate into macrophages.

In contrast to MMϕ isolated at P10, MMϕ at P56 were characterised by the emergence of differentiating MMФ and NA-MMФ. NA-MMФ were characterised by the upregulation of canonical microglia genes, such as *Tmem119, Olfml3* and *Hexb* (Figure 2F, S2H-I). Of interest, this cluster highly expressed the tight junction protein F11r (JAM-A), typically expressed by endothelial cells in the gastrointestinal tract^25^. Indeed, F11r was expressed by endothelial cells within the muscularis externa at both P10 and P56 (Figure 3F). However, at P56 but not at P10, a subset of MMФ expressing F11r^+^ could be observed in close proximity to neuronal cell bodies in the myenteric plexus, and to neuronal fibres within the muscle layer (Figure 3F). Quantification via flow cytometry revealed that F11r^hi^ NA-MMϕ are present almost exclusively at P56, in line with our scRNAseq findings (Figure S2B,3G).

Of note, the most abundant cluster at P56 was composed of differentiating MMФ. We did not identify surface marker expression restricted to differentiating macrophages, and these cells were thus identified as Lyve1^-^ F11r^-^ Ki67^-^ MMФ (Figure S2B). Lyve1^-^ F11r^-^ Ki67^-^ MMФ were more abundant at P56 compared to P10 mice, in line with our scRNAseq analysis (Figure 3H). To understand whether these cells may ultimately differentiate into one of the identified mature clusters, we performed an RNA Velocity analysis on all non-replicating cells to estimate the direction of transcriptional shifts^26^. This analysis indicated a potential differentiation trajectory from differentiating MMϕ to NA-MMϕ, suggesting that these cells may ultimately differentiate into NA-MMϕ (Figure 3I).

Finally, to determine whether the frequency of the identified clusters undergoes further age-related shifts, we compared the cluster distribution in 8 and 20 weeks old mice using flow cytometry. Differentiating Lyve1^-^ F11r^-^ Ki67^-^ MMФ decreased in frequency, while the abundance of F11r^hi^ NA-MMФ increased with age (Figure 3K). Interestingly, the frequency of Lyve1^+^ MMϕ also decreased with age, representing less than 5% of all MMϕ at 20 weeks of age compared to 14% at 8 weeks of age (Figure 3K). Taken together, we identified distinct subpopulations of MMϕ that change their abundance during development and age and occupy specific subtissular niches in the muscularis.

### ENS-derived TGFβ Instructs MMϕ to Adopt a Neuron-Associated Phenotype

It is well established that the niche in which tissue resident macrophages reside largely determines their phenotype and transcriptome^11^. We next aimed to investigate the niche-driven imprinting of the neuron-associated phenotype in MMϕ after weaning, and which molecular mediators may be involved in shaping this phenotype. Transforming Growth Factor β (TGFβ) is a signalling molecule that is encoded by 3 isoforms in mammals, TGFβ1, 2 and 3. In the brain, TGFβ2 has been shown to be crucial in conferring the mature microglia signature^28^. As NA-MMϕ upregulate numerous microglia signature genes, we sought to investigate a possible role of TGFβ in imprinting the NA-MMϕ phenotype. Thus, the expression of TGFβ1, 2 and 3 was assessed in the muscularis externa at P10 and at P56. Interestingly, TGFβ3 expression increased significantly at P56 compared to P10 (Figure 4A). To determine whether TGFβ may drive the expression of genes expressed by NA-MMϕ, bone marrow-derived macrophages (BMDMs) were cultured in the presence of TGFβ1, 2 or 3. Incubation with TGFβ led to the upregulation of canonical NA-MMϕ genes, including *Hexb, F11r, Cx3cr1* and *Ccrl2* (Figure 4B, S4A). However, some genes upregulated in NA-MMФ were not upregulated by TGFβ, including *Tmem119* and *Olfml3*, suggesting that additional molecular mediators may be involved to imprint the full NA-MMϕ phenotype (figure S4A).

**Figure 4:**
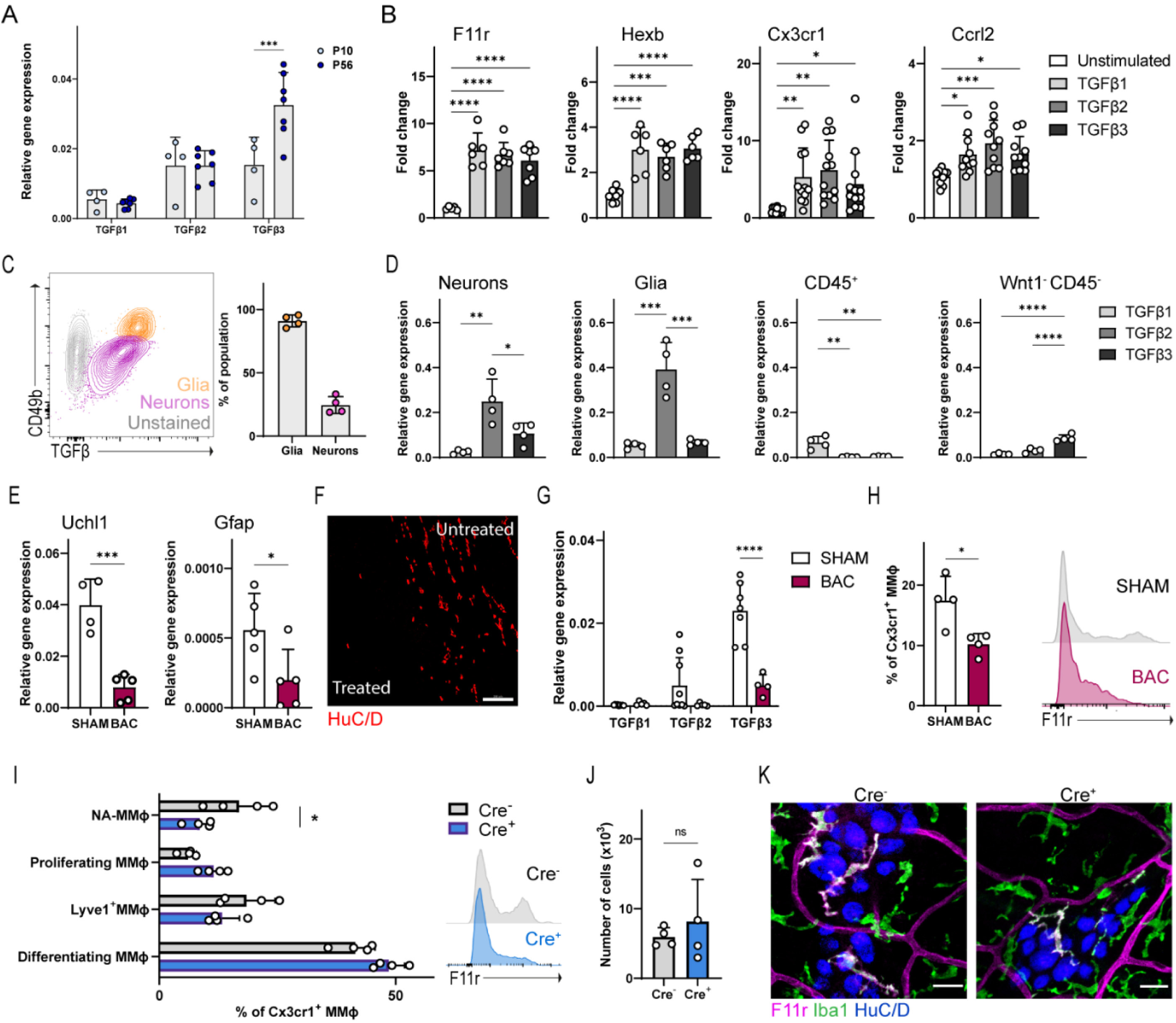
ENS-derived TGFβ instructs the NA-MMϕ phenotype. (**A**) Gene expression of TGFβ orthologues within the muscularis externa at P10 and P56. (**B**) Gene expression of NA-MMϕ marker genes in BMDM following 24h stimulation with TGFβ1, TGFβ2 or TGFβ3. (**C**) Representative contour plot of TGFβ expression in enteric neurons and enteric glia, as assessed via flow cytometry. Cells were pre-gated on live CD45^-^ Wnt1^+^. The graph depicts the percentage of neurons and glia expressing TGFβ. (**D**) Gene expression of TGFβ1, TGFβ2 and TGFβ3 in enteric neurons (Wnt1^-^Cd49b^-^), enteric glia (Wnt1^+^Cd49b^+^), immune cells (CD45^+^) and non-immune non neural (Wnt1^-^ Cd49b^-^) cells sorted from the muscularis externa of Wnt1^cre/wt^ Rosa26^tdT/WT^ mice. (**E**) Gene expression of Uchl1 and Gfap in the muscularis externa 5 days after enteric denervation with BAC. (**F**) Representative confocal image depicting BAC treated and untreated muscularis externa. Scale bar is 200µm. (**G**) Gene expression of TGFβ1, TGFβ2 and TGFβ3 in the muscularis externa 5 days after enteric denervation with BAC compared to SHAM. (**H**) Percentage of F11r^hi^ Cx3cr1^+^ macrophages in mice denervated with BAC compared to controls (SHAM). Representative histogram of F11r expression in live CD45^+^ Cd11b^+^ CD64^+^ Cx3cr1^+^ cells. (**I**) Frequency of MMФ populations in the muscularis externa of Cx3cr1^creERT2/WT^ TGFbr2^fl/fl^ (Cre^+^) and Cx3cr1^WT/WT^ TGFbr2^fl/fl^ (Cre^-^) mice, as assessed via flow cytometry. Representative histogram showing F11r expression in Cd11b^+^ CD64^+^ Cx3cr1^+^ cells of Cx3cr1^creERT2/WT^ TGFbr2^fl/fl^ (Cre^+^) and Cx3cr1^WT/WT^ TGFbr2^fl/fl^ (Cre^-^) mice, as assessed via flow cytometry. (**J**) Number of live Cx3cr1^+^ Cd64^+^ Cd11b^+^ Cd45^+^ macrophages in the muscularis externa of Cx3cr1^creERT2/WT^ TGFbr2^fl/fl^ (Cre^+^) and Cx3cr1^WT/WT^ TGFbr2^fl/fl^ (Cre^-^) mice as assessed via flow cytometry. (**K**) Representative confocal image of the myenteric plexus of Cx3cr1^creERT2/WT^ TGFbr2^fl/fl^ (Cre^+^) and Cx3cr1^WT/WT^ TGFbr2^fl/fl^(Cre^-^) mice. Scale bar is 20µm. Data are shown as Mean ± SD, and are representative or pooled from at least 2 independent experiments. Data are analysed using T-tests or ANOVA followed by Holm-Sidak multiple comparisons. * *p* < 0.05; ** *p* < 0.01;*** *p* < 0.001; **** *p* < 0.0001; ns = not significant.

To determine the source of TGFβ in the muscularis externa, expression of TGFβ was assessed via flow cytometry using a non-isoform specific antibody. As NA-MMϕ are located in close proximity to the ENS, we hypothesised that enteric neurons or glia may represent the source of TGFβ. To assess TGFβ expression in the ENS, transgenic mice expressing tdTomato in neural crest-derived cells (*Wnt1*^*cre/wt*^*Rosa26*^*tdT/wt*^) were used. As this genetic construct labels both enteric neurons and enteric glia, enteric glia were distinguished from enteric neurons by CD49b expression, as previously described^29^. TGFβ expression was detected within the tdTomato^+^ population, with almost all enteric glia expressing TGFβ and around 25% of enteric neurons also expressing TGFβ (Figure 4C). Of note, TGFβ was also detected in Wnt1^-^ cells, suggesting that there may be other sources of TGFβ within the muscularis externa (Figure S4B). We did not detect TGFβ expression within the immune cell compartment (Figure S4B). To further validate these findings, enteric neurons (Wnt1^+^CD49b^-^), enteric glia (Wnt1^+^CD49b^+^), immune cells (CD45^+^) and non-neural CD45^-^ cells were sorted from the muscularis externa and analysed for expression of TGFβ1, 2 and 3 via qRT-PCR. TGFβ1 expression was detected only at very low levels within all sorted populations (Figure 4D). TGFβ2 was expressed by enteric glia and enteric neurons, while TGFβ3 was mainly detected in enteric neurons and non-neural CD45^-^ cells (Figure 4D). These findings confirm the cells of the adult ENS serve as a source of TGFβ within the muscularis externa.

Finally, to further confirm the importance of ENS-derived TGFβ in determining the NA-MMϕ phenotype, we performed ENS ablation in the muscularis externa using benzalkonium chloride (BAC), a cationic surfactant which, when applied directly to the muscularis externa, causes extensive neurodegeneration^30,31^. BAC treatment led to ablation of the ENS in the treated portion of the muscularis externa 5 days after treatment, as shown by the reduced expression of neuronal (*Uchl1*) and glial (*Gfap*) gene expression, and the loss of HuC/D^+^ neurons in the myenteric plexus (Figure 4E,F). BAC treatment led to a significant reduction of TGFβ3 expression only, whereas TGFβ2 expression remained unchanged (Figure 4G). To assess the role of neuron-derived TGFβ on MMϕ phenotype, we assessed NA-MMϕ via flow cytometry 14 days after BAC denervation. Loss of the ENS induced a reduction of the F11r^hi^ population within the MMϕ compartment, while global MMϕ cell counts were not affected (Figure 4H, S4C). Finally, we assessed the MMϕ compartment in *Cx3cr1*^*CreERT2*^ *Tgfbr2*^*fl/fl*^ mice, in which the TGFβ receptor type II, Tgfbr2, is excised from all Cx3cr1-expressing macrophages upon tamoxifen administration. TGFβ-signalling requires binding of TGFβ to Tgfbr2, that may then phosphorylate the TGFβ receptor type I (Tgfbr1), enabling the kinase activity of Tgfbr1^32,33^. *Cx3cr1*^*CreERT2*^ *Tgfbr2*^*fl/fl*^ mice were thus treated with tamoxifen after weaning (at 5 weeks of age) and MMϕ populations were assessed via flow cytometry at 10-12 weeks of age. Loss of TGFβ receptor signalling led to a significant reduction in NA-MMϕ in Cre+ mice compared to Cre-controls, while other MMϕ populations and global MMϕ cell counts were not affected (Figure 4I-K). Taken together, these findings suggest that ENS-derived TGFβ imprints the NA-MMϕ phenotype observed in the adult muscularis externa.

## Discussion

Correct development and maturation of the ENS is critical for survival. Especially early in life the gastro-intestinal tract, and thus also the ENS, has to significantly adapt its function to the type of food ingested, changing from milk to solid food at the time of weaning. In the here presented study, we present a novel role of MMϕ in the postnatal maturation of the ENS. Furthermore, we describe different MMϕ sub-populations which change in their composition with age and serve distinct functions in sub-anatomical niches in the muscularis externa, including a distinct, neuron-associated MMϕ subset which expresses a microglia-like phenotype. Of note, this microglia-like phenotype is instructed by the ENS itself, via production of TGFβ. Our findings indicate that the ENS, similar to the brain, is shaped and maintained by a dedicated population of tissue resident macrophages, that adapts its phenotype and transcriptome to the timely needs of the ENS niche.

### Remodelling of the ENS by MMϕ

Although the intestine is colonized by enteric neurons at birth, the ENS still requires significant remodelling before it acquires its typical organization into ganglia connected by nerve tracks^6,7^. Reorganisation of the ENS in early postnatal life requires refinement of enteric neurons, the number of ganglia and the fibers that connect the ganglia to each other. Here, we show that depletion of MMϕ during this critical refinement period led to hyperganglionosis and a disorganised immature ENS structure, demonstrating that MMϕ are responsible for the reorganisation of the ENS during early postnatal life. Moreover, our findings are in line with previous studies using *Csf1*^op/op^ mice, which are devoid of MMϕ, and which display a disorganised ENS and increased neuronal density in adulthood^34^. In line with these findings, when we performed transcriptional profiling of MMϕ at P10, we observed that MMϕ at P10 were predominantly Lyve1^+^ MMϕ. Lyve1^+^ MMϕ upregulated numerous canonical phagocytosis genes, and presented superior phagocytic capacity as tested in an *ex-vivo* phagocytosis assay compared to other identified MMϕ subpopulations. Taken together, these findings demonstrate that early in life, MMϕ play a crucial role in remodelling the ENS, enabling it to adapt to the evolving needs of the gut as it transitions to solid food.

### ENS-MMϕ Relationship in Adulthood

We previously demonstrated that loss of MMϕ in adulthood leads to significant neurodegeneration and impaired GI function^17^. Here we confirmed these findings; depletion of MMϕ at 8 weeks of age led to a significant reduction in neuronal density, suggesting that in adulthood, MMϕ shift toward a neuro-supportive function. In line, scRNAseq of P56 MMϕ revealed the emergence of a novel, neuron-associated MMϕ (NA-MMϕ) population. This NA-MMϕ population upregulated genes including *Tmem119, Hexb, Olfml3* and their surface marker *F11r*, and were found to be associated to neuronal cell bodies and fibres in mice in adulthood. As we previously demonstrated that MMϕ preferentially associated to enteric neurons are long-lived, we speculate that this population represents a specialised population of MMϕ that is critical for neuronal support and survival in adulthood^17^.

### The ENS-MMФ Niche

Recently, we described a macrophage population associated to neurons in the enteric nervous system, which displayed an upregulation of canonical microglia genes^17^. Since then, several similar ‘microglia-like’ macrophage populations have been reported to be associated to large peripheral nerve bundles including dorsal root ganglia, vagal nerve, subcutaneous fascial nerves and sciatic nerves^22,23^. The definition of ‘microglia-like’ largely derives from the transcriptional signature, as these cells have been shown to upregulate canonical microglia genes, including *Hexb, Siglech, Trem2* and *Olfml3*^35,36^. It should be emphasized though that they fail to upregulate the typical microglial transcription factor Sall1, suggesting that distinct anatomical niches might still vary to a certain amount in the factors released to specify nerve-associated macrophages^22,23,35^. Intriguingly, while neuron-associated macrophage subsets share a microglia-like transcriptional signature, the signals inducing this signature have not been identified. Here, we identify TGFβ, a cytokine critical for microglial instruction in the brain, as a molecular cue employed by the ENS to instruct local macrophages to adopt a neuron-associated profile. *In vitro* stimulation of BMDM with TGFβ led to the upregulation of genes identified in NA-MMϕ while *in vivo*, chemical ablation of the ENS or genetic inhibition of TGFβR signalling in MMϕ led to the reduction of NA-MMϕ in the muscularis externa. TGFβ-TGFβR signalling is thus crucial for the maintenance of NA-MMϕ in the adult muscularis externa. This is, to our knowledge, the first study which identifies the molecular mediator that defines the neuron-associated macrophage niche. To what extent TGFβ might be a conserved signal used by neuronal niches throughout the body to imprint a specific phenotype to the local macrophage pool requires further study.

### Functional Plasticity of MMФ

With the notable exception of microglia, the notion that macrophages within tissues retain the plasticity to adapt their function in order to respond to evolving needs of the tissue has largely been restricted to the context of disease and inflammation. Here, we demonstrate that resident MMϕ undergo profound transcriptional maturation from postnatal development to adulthood, a maturation that is likely cued by the surrounding cells and the evolving needs of the tissue. A similar developmental plasticity has been demonstrated in the brain, where microglia are critical for the refinement of synaptic connectivity during early postnatal development and for neuronal homeostasis and neurogenesis in adulthood^37–39^. Also in the brain, this functional adaptation is mirrored by transcriptional shifts that enable microglia to respond to the evolving requirements of the rapidly developing surrounding cells, which is crucial for the development and maturation of the central nervous system (CNS)^40^. Strikingly, the global transcriptional shifts observed in MMϕ resemble those observed in microglia, with a global upregulation of genes involved in phagocytosis and replication during early postnatal life, and subsequently, in adulthood, upregulation of genes involved in immune surveillance^40^. Similarly, changes in morphology and proliferative rates of MMϕ also paralleled microglial differentiation^41,42^.

### Paralleled Role for Macrophages/Microglia in the Central and Enteric Nervous System

The finding that macrophages participate in the process of neuronal refinement in the ENS opens up new horizons in our understanding of neurodevelopmental disorders of the ENS. In the brain, depletion of microglia during a time window critical for synaptic pruning leads to altered learning and memory, social, mood-related, and locomotor behaviours^43,44^. Of note, hyperganglionosis of the myenteric and submucous plexus is associated to severe constipation in children, and has been associated with gastrointestinal motility disorders^2,3^. Whether these abnormalities in ENS morphology and gut dysfunction can be ascribed to insufficient MMϕ function early in life requires further study. Conversely, based on our findings, MMϕ dysfunction in adulthood should be considered as a potential disease mechanism in neurodegeneration in the ENS, as described during aging and in diseases such as diabetes and obesity^17,45,46^. The resemblance between the neuronal niche in the muscularis externa and the brain, where TGFβ is critical for the instruction of microglia, is striking, and further supports the notion that the ENS may resemble the CNS more than previously anticipated. Indeed, it is widely reported that neurodegenerative and neurological disorders of the CNS, including Parkinson’s disease, Alzheimer’s disease and Multiple Sclerosis, present wide-ranging ENS symptomatology^4,47,48^. It is intriguing to speculate that the ENS, as the ‘brain of the gut’, may mirror the CNS in homeostasis and in disease, and may thus offer an unexploited snapshot of ongoing pathological processes in the brain.

## Acknowledgements

We thank Pier Andrée Penttila and Reena Chinnaraj from the KU Leuven FACS core for their excellent support and cell sorting. We also thank the Cell Imaging Core (Pieter Vanden Berghe and Tobie Martens, KU Leuven) for confocal imaging (supported by AKUL 15/37 and FWO I001918N). MFV is supported by FWO PhD fellowship 11C2219N. GB is funded by ERC Advanced grant number 833816-NEUMACS.

## Author Contributions

MFV designed experiments, conducted experiments and wrote the manuscript. MCP performed histological analyses. EM and AS performed analysis of scRNAseq data. IA and NF provided excellent technical support throughout all experiments. MD, NS, TM, KV, JVH performed experiments. SV, SDS, PVB,TV and AS provided intellectual input. KK and MP provided mice and PP performed genotyping and tamoxifen injections. GB led the project and revised the manuscript.

## Materials and Methods

### Mice

All animal experiments were performed in accordance with the European Community Council and approved by the Animal Care and Animal Experiments Committee of the Medical Faculty of the KU Leuven. Mice were maintained under a 14h/10h dark/light cycle, at a temperature of 20-22 °C, provided with food and water ad libitum. Both male and female mice were used for this study. *Cx3cr1*^*GFP/GFP*^ mice were obtained from Steffen Jung, Weizmann Institute of Science^49^. Wnt1.Cre mice were obtained from Pieter Vanden Berghe^50^. *Rosa26*^*TdT/TdT*^ mice were obtained from Jackson Laboratory. *Wnt1*^*Cre/WT*^ *Rosa26*^*TdT/WT*^ *Cx3cr1*^*GFP/WT*^ mice were bred in our facility and maintained on a C57Bl/6 background. *Cx3cr1*^*CreERT2/WT*^ *Tgfbr2*^*fl/fl*^ mice were kindly provided by Katrin Kierdorf and Marco Prinz, University of Freiburg.

### Histology and immunofluorescence

The ileum was dissected, carefully flushed using Hank’s Balanced Salt Solution (HBSS) (Gibco) supplemented with fetal calf serum (FCS, 3%, Lonza) and HEPES (1.8%) and pinned on a sylgard plate. The tissue was cleaned of mesenteric fat, opened longitudinally and pinned with the muscle layer facing upwards. The muscle layer was carefully peeled from the submucosa, pinned flat on the sylgard plate and then fixed for 30 minutes in PFA 4%. Fresh whole mount tissue was stained within one week of collection. Tissue was permeabilized in PBS supplemented with Triton X-100 (0.3%; permeabilization buffer) for 2 hours at room temperature, and then blocked with permeabilization buffer supplemented with Donkey serum (5%) and Bovine serum albumin (BSA, 5%; blocking buffer) for 2 hours at room temperature. Tissue was incubated with primary antibodies diluted in blocking buffer overnight at 4°C. Tissue was extensively washed in PBS prior to incubation with secondary antibodies diluted in blocking buffer for 2 hours at room temperature. Samples were washed and mounted using SlowFade Diamond Antifade mounting medium (Thermofischer Scientific) and imaged using a Zeiss LSM880 confocal microscope. Negative controls (incubated with secondary antibody, but no primary antibody) were included for every analysis. All antibodies used are listed in supplementary table 1.

### Quantification of synaptic volume and synaptic engulfment

Quantification of synaptic engulfment was carried out using a published protocol adapted to our purpose^20^. Briefly, Z-stacks were imported to Imaris (Bitplane) for analysis, where digital 3D reconstructions were performed to calculate MMϕ volume, and synaptic machinery. Engulfment was determined to be the volume of synaptic machinery engulfed by MMϕ, normalised to the volume of the MMϕ. Synaptic density was determined as the volume of synaptic machinery, normalised to the volume of the region of interest.

### *Ex-vivo* live imaging

The ileum of *Wnt1*^*Cre/WT*^ *Rosa26*^*TdT/WT*^ *Cx3cr1*^*GFP/WT*^ mice was carefully in carbogenated Krebs solution at 4°C. The mucosal, submucosal, and longitudinal muscle layers were carefully removed to obtain a circular muscle with adherent myenteric plexus preparation which was mounted over a small inox ring, immobilized by a matched rubber O-ring^51^. Inox rings with muscularis externa preparations were positioned in FluoroDishes and maintained at 37°C with 5% CO2 in Krebs solution with 1µm nifedipine for time-lapsed imaging using a LSM880 Airyscan confocal microscope (Zeiss). Stitched videos were analysed and processed using ImageJ software (Fiji).

### Analysis of MMϕ morphology

Z-stacks of confocal images were imported into Imaris software (Bitplane). The filament tool was used to trace the filopodia of each MMϕ manually. Number, length and branching of filopodia was calculated using the filament tool. MMϕ cell volume was estimated using the volume tool.

### Tissue harvest for single cell suspensions

The small intestine was removed from mesenteric fat tissue, opened longitudinally and cleaned in Hank’s Balanced Salt Solution (HBSS) (Gibco) supplemented with fetal calf serum (FCS, 3%, Lonza) and HEPES (1.8%). The opened tissue was pinned to a sylgard plate and the muscle layer was carefully dissected and collected in RPMI supplemented with FCS (3%) and HEPES (2%). The tissue was cut in small pieces and incubated with Collagenase IV (500U/ml) and DNAse for 30 minutes at 37°C in constant agitation. After digestion, cells were filtered through a 70 µm cell strainer and washed using PBS supplemented with FCS (2%) and EDTA (1.6mM) and counted prior to staining for Flow Cytometry.

### Flow cytometry

Single cell suspensions were blocked using anti CD16/CD32 (Fc block) and subsequently labelled for flow cytometry analysis using fluorophore-conjugated anti-mouse antibodies for 30 min at 4°C using the antibodies listed in supplementary table 2. For intracellular staining, cells were fixed using the FoxP3 Intracellular Staining kit (eBioscience), and incubated with conjugated intracellular antibodies for 1 hour at 4°C. Samples were acquired using a Symphony A5 (BD Biosciences). Analysis was carried out using FlowJo (v10, Tree Star).

### Depletion of muscularis macrophages

Muscularis macrophages were depleted as previously described^52^. Briefly, mice received 2 injections (37.5 µg/g) of either α-CSF1R (Clone AFS98, Bio X Cell) or IgG2a isotype control (Clone 2A3, Bio X Cell) I.P., 3 days apart, and were euthanized via CO2 overdose 4 days after the last injection.

### scRNAseq library preparation

Single-cell suspensions were obtained from the ileum of Cx3cr1^GFP/WT^ mice as previously described. Cells were counterstained with DAPI to exclude dead cells and sorted using an Aria III cell sorter (BD) into skirted 96-well plates for scRNAseq. cDNA libraries from 376 cells were generated based on the Smart-seq2 protocol^53^. Briefly, mRNA was reverse transcribed and cDNA was amplified via PCR. Amplification was carried out using KAPA HIFI Hot Start ReadyMix (Roche, 07958919001) and purification by magnetic beads (CleanNA CPCR). Quantity and quality of cDNA was assessed with the quantifluor RNA system (Promega E3310) and Agilent 2100 BioAnalyzer with a high-sensitivity chip. Library was prepared using the Nextera XT library prep and index kit (Illumina). cDNA was tagmented by transposase Tn5 and amplified with dual-index primers. Reagents were mixed together by the Echo 555 (Labcyte) and pooled-nextera XT libraries were purified. Single-cell libraries were pooled together and sequenced single-end 50 bp on a single lane of a HiSeq4000 (Illumina). All results related to scRNA-seq are based on 4 pooled biological replicates per group.

### scRNAseq analysis

Raw fastq reads were obtained from the sequencer and underwent a series of steps before being imported to Seurat (v4.1.0) for analysis. The reads were trimmed with cutadapt 3.0 to remove adapters. Reads were then mapped to the mouse genome (GRCm38.p6) with STAR 2.7.6a and reads that mapped to a gene were counted with htse-count 0.12.4. Metrics were then collected and optical duplicates were marked using Picard 2.23.8. Finally, multiqc 1.9 was used to collect all the analyses log files into a single report and countmatrix files were generated (genes per cell counts). After pre-processing of the reads, each library was matched with a sample using the predetermined Nextera barcodes. Following a quality check and filtering of these samples (features: 2000-7500; UMIs > 2000; mitochondrial gene content < 5%), cells were normalized, scaled and batch-corrected (5000 anchors and using 35 neighbors to filter the anchors) using Seurat. Highly variable genes (i.e. 1742) were then used for PCA dimensionality reduction and visualized using 2-D Uniform Manifold Approximation and Projection (UMAP) in Seurat. Cells were grouped by unsupervised clustering at different clustering resolutions and the clustering resolution 1.1 was chosen to reflect clear biologically-driven transcriptional differences between clusters. The smallest cluster of the dataset, comprising of 34 cells, was excluded from further analysis due to cargo contamination (Figure S2D-G). All further analyses, including identification of marker genes, single-cell regulatory network inference and clustering (SCENIC) and RNA Velocity were performed after removal of this cluster.

Gene regulatory network analysis was performed using pySCENIC (v.0.9.3) using default parameters. SCENIC is a tool to infer gene regulatory networks from single-cell RNA-seq data^27^. Differentially activated regulons were determined by performing a two-sided Kolmogorov–Smirnov non-parametric rank sum test on the regulon AUC values of cells in the various clusters, P values were false discovery rate (FDR)-adjusted using the Benjamini–Hochberg method and regulons with an adjusted P value less than 0.01 were considered differentially activated.

To perform lineage trajectory analysis, we employed RNA Velocity. RNA Velocity is a high-dimensional vector that predicts the future state of individual cells on a timescale of hours Splicing information of reads (spliced/unspliced) was extracted from the binary file containing the sequence aligned data (.bam) from a 10X run (10X genomics) of each separate sample using the velocyto (01.17.17) pipeline (http://velocyto.org). It was then stored in a loom file and then read into an AnnData with python (v 2.7.5) object for downstream analysis with scVelo (0.2.4) and numpy (1.19.4). For the estimation of velocities and the RNA force field the recommended steps of scVelo were followed, namely: Filtering, normalisation and log transformation of the data, computing first- and second-order moments, estimating velocities, and constructing a velocity graph. First- and second-order moments were calculated for each cell across its nearest neighbors of a single-cell neighbourhood-graph in PC space (number of neighbors=30, number of PCs=35). Then estimates for velocities were obtained by fitting a dynamic model of transcription for each gene. Finally, a velocity graph was computed from the correlations between potential cell transitions in the neighbourhood graph and the predicted cell state change given by the velocity vector. This graph was then used to project the estimated velocities into the original low dimensional UMAP space.

### Ex-vivo phagocytosis assay

Cells were isolated to obtain single-cell suspensions. To assess ex-vivo phagocytosis, cells were incubated for 45 minutes at 37°C in the presence of Texas Red-conjugated Dextran (MW 70 000, Invitrogen) prior to extracellular surface marker staining, while controls were maintained at 4° in the presence of Texas Red-conjugated Dextran. Cells were further processed as described for flow cytometry. Phagocytic index is defined as percentage of cells engulfing dextran, multiplied by the MFI (mean fluorescence intensity) of the fluorescently labelled dextran engulfed by the cells.

### RNA Isolation and qRT-PCR

RNA from tissue or cells was extracted using the innuPREP RNA Mini Kit 2.0 (AnalytikJena), according to manufacturers’ guidelines. RNA was retrotranscribed using the qScript cDNA Synthesis Kit (Quantabio) according to manufacturers’ instructions using a GeneAmp PCR System 2700 (Applied Biosystems). qRT-PCR was performed using LightCycler® 480 SYBR Green I master mix (Lifescience, Roche) on a Lightcycler 480 (Roche) using the primers detailed in supplementary table 3. All data were normalized to Rpl32 quantified in parallel amplification reactions.

### BMDM experiments

Femura bones from 8 week old C57/Bl6J mice were isolated and collected in RPMI. The extremities of the bones were cut, and a 24G gauge needle was used to flush out the bone marrow into complete RPMI (10% FCS, 1% P/S, 1% L-glutamine, 1% Sodium Pyruvate and 1% Hepes). The cells were then centrifuged, resuspended in red cell lysis buffer for 1 minute, then washed with PBS and centrifuged. The pelleted cells were then resuspended and plated in complete RPMI supplemented with 7.5% L929 medium. Cells were grown for 7 days prior to stimulation with either 2ng/ml of TGFb1, 2 or 3. 24 hours after stimulation, cells were collected for RNA extraction and qRT-PCR analysis.

### Sorting of live cells for gene expression analysis

Single-cell suspensions were obtained from the ileum of Wnt1^Cre/WT^ Rosa26^TdT/WT^ Cx3cr1^GFP/WT^ mice as previously described, and further processed for staining with cell-surface markers. Cells were sorted using a Sony MA900 cell sorter directly into RLT buffer additioned with β-mercaptoethanol. After sorting, samples were vortexed as per manufacturer’s instructions and stored at −80°C until further analysis.

### Enteric denervation using Benzalkonium Chloride (BAC)

Animals were anesthetized by i.p. injection of ketamine (Ketalar; 100 mg/kg; Pfizer) and xylazine (Rompun; 10 mg/kg; Bayer). A laparotomy with a midline incision was performed, and a portion of the ileum was exposed. Sterile cellulose filter paper disks (5-mm diameter; Whatman) soaked in 0.1% BAC (Sigma-Aldrich) in NaCl solution were placed on the exteriorized intestine, 2 cm from the ileocecal valve. Sham mice received the same procedure, however cellulose filter disks were soaked in NaCl solution online. The tissue was incubated for 15 min with BAC or NaCl, after which the paper disk was removed and the treated area thoroughly rinsed with sterile saline solution. Mice were killed with CO2 overdose 14 days after treatment. The treated segments of bowel were harvested from each animal, and the muscularis layer was isolated for flow cytometry analysis as previously described.

### Tamoxifen treatment

For activation of Cre recombinase, tamoxifen (TAM, Sigma-Aldrich, Taufkirchen, Germany), was dissolved in sterile corn oil (Sigma-Aldrich, Taufkirchen, Germany) prior to injection at a concentration of 20mg/ml. Cre recombination was induced by injecting 4 mg TAM/200μl corn oil subcutaneously twice 48 h apart.

### Statistics

Statistical analysis was performed using Prism 9 (GraphPad Software). Data are represented as Mean ± SD and analysed using T-tests, ANOVA (followed by Tukey’s multiple comparisons) or Kruskal-Wallis (followed by Dunn’s multiple comparisons). Statistical outliers were removed using Grubbs outlier test. A *p* value < 0.05 was considered statistically significant.

