## Supplementary Information for "Neuro-immune Crosstalk in the Enteric Nervous System from Early Postnatal Development to Adulthood"

**Supplementary Material**

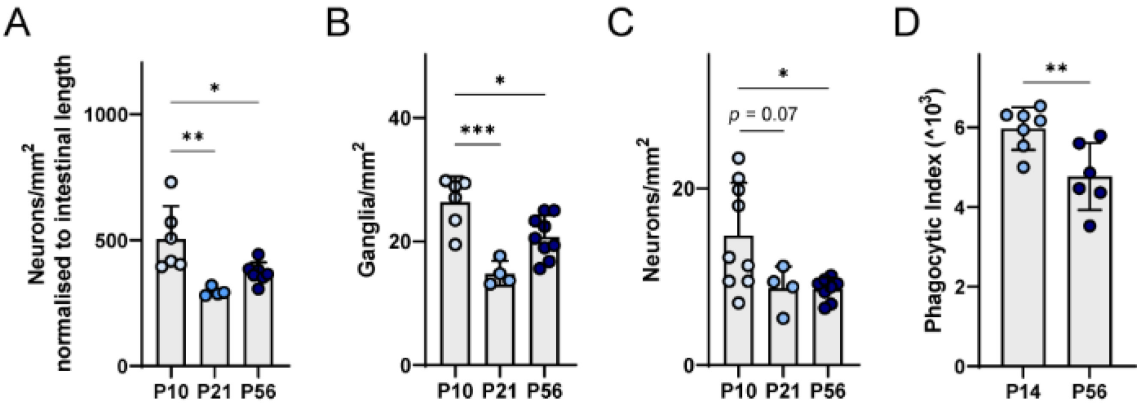


Figure S1: Quantification of neuronal density (**A**), the number of ganglia (defined as containing 5 or more neurons) per mm$^2$ (**B**), and the number of single extra-ganglionic neurons per mm$^2$ (**C**), in the myenteric plexus at 10 (P10), 21 (P21) and 56 (P56) days of age, corrected for tissue growth. (**D**) Mean fluorescence intensity of MMϕ engulfing neural cells at 14 days (P14) and 56 days (P56) of age. Unpaired t-tests or Ordinary One-Way ANOVAs with Tukey's multiple comparisons. Data are shown as Mean ± SD, and are representative of at least 2 independent experiments. * $p < 0.05$; ** $p < 0.01$; *** $p < 0.001$.

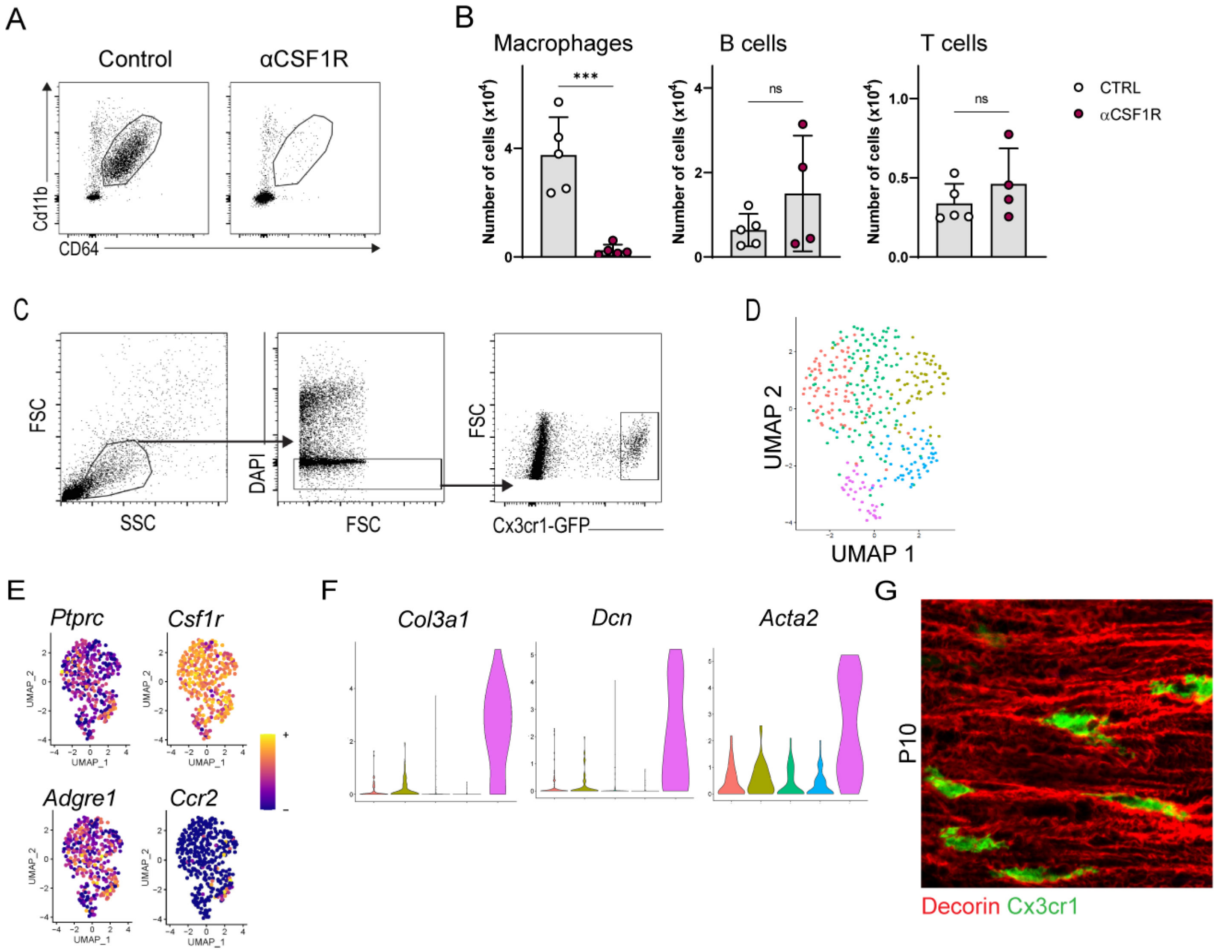


Figure S2: (**A**) Representative dot plot in αCSF1R-treated mice and controls to show efficient depletion of MMϕ. Cells shown are pre-gated on fsc/ssc, single cells, live and $CD45^+$. (**B**) Quantification of immune cells in αCSF1R-treated mice and controls. Macrophages are defined as $CD45^+CD11b^+CD64^+$, B cells as $CD45^+CD3e^-CD64^-CD11b^-CD19^+$ and T cells as $CD45^+CD3e^+$. All populations were pre-gated on fsc/ssc and live cells. Unpaired t-tests. Data are shown as Mean ± SD. *** $p < 0.001$; ns = not significant. (**C**) Gating strategy used to isolate $Cx3cr1^+$ MMϕ from the muscularis externa for scRNAseq. (**D**) UMAP of data prior to exclusion of contaminating cluster (pink). (**E**) UMAP with heatmap colour-coding to show expression for canonical macrophage markers and CCR2 within the scRNAseq dataset. (**F**) Violin plots showing the upregulation of extracellular matrix genes in the contaminating cluster (pink). (**G**) Representative confocal image of the muscularis externa at P10, showing $Cx3cr1^+$ MMϕ embedded within the extracellular matrix (Dcn, Decorin).

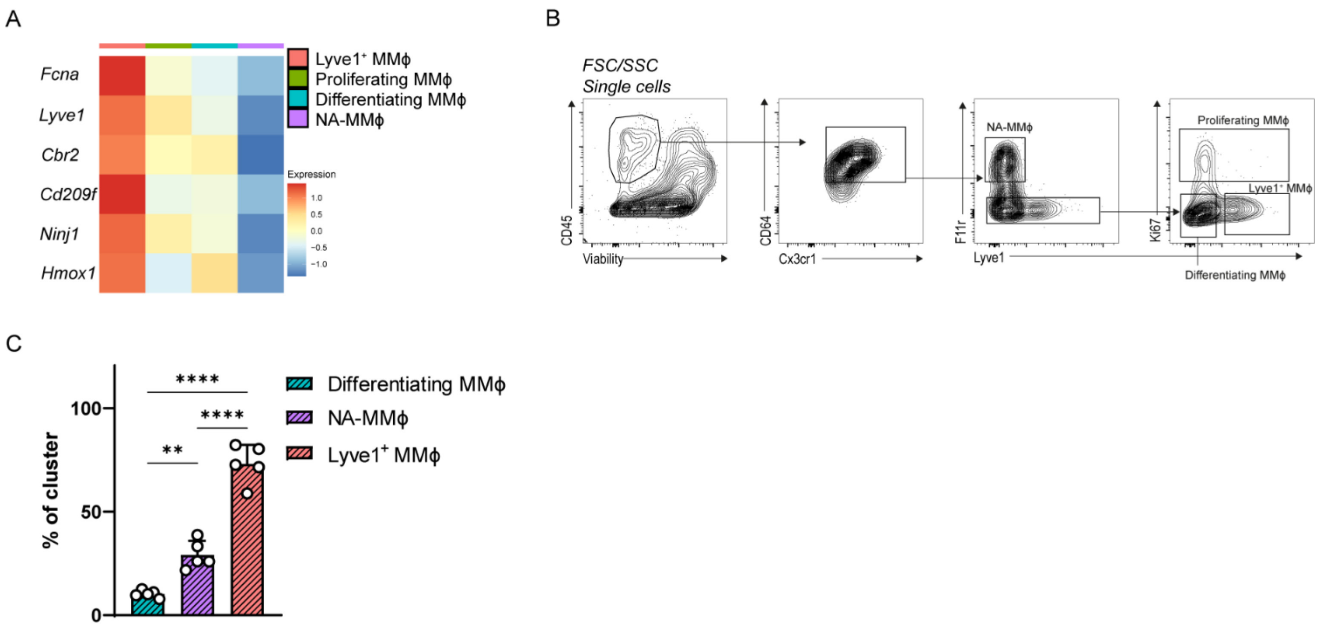


Figure S3: (**A**) Heatmap depicting expression of genes identified by Chakarov et al., within each cluster, cells pooled. (**B**) Representative gating strategy used to identified MMϕ subsets via flow cytometry. (**C**) Percentage of cells of each MMϕ subset phagocytosing fluorescently labelled dextran. Data are shown as Mean ± SD, analysed ANOVA followed by Tukey's multiple comparisons. ** $p < 0.01$; **** $p < 0.0001$.

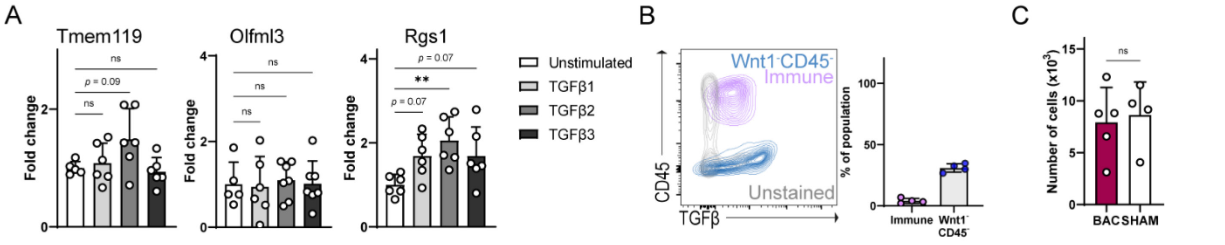


Figure S4: (**A**) Gene expression of NA-MMϕ marker genes in BMDM following 24h stimulation with TGFβ1, TGFβ2 or TGFβ3. (**B**) Representative contour plot showing TGFβ expression in immune cells (CD45$^+$) and Wnt1$^-$CD45$^-$ cells. Only live cells are shown. The graph depicts the percentage of cells of each population expressing TGFβ. (**C**) Number of live CD45$^+$ CD64$^+$ CD11b$^+$ Cx3cr1$^+$ MMϕ detected in the muscularis externa in SHAM and BAC-treated mice, normalised to cell count. Data are shown as Mean ± SD, analysed using T-test or ANOVA followed by Holm-Sidak multiple comparisons. ** $p < 0.01$; ns = not significant.

Supplementary table 1: Antibodies used for immunofluorescence

| Antibody | Source | Cat. Number |
|---|---|---|
| Anti-GFP | Nacalai Tesque | 04404-26 |
| Anti-HuC/D | Supplied by V. Lennon (Mayo Clinic) | |
| Anti-Neurofilament | Abcam | ab72996 |
| Anti-Synapsin I | Abcam | ab64581 |
| Anti-Decorin | R&D Systems | AF1060-S |
| Anti-Lyve1 | Abcam | ab14917 |
| Anti-VE-Cadherin | R&D Systems | AF-1002 |
| Anti-F11r (JAM-A) | R&D Systems | AF-1077 |
| Anti-Ki67 | Abcam | ab15580 |
| Anti-βIII-Tubulin | Abcam | ab78078 |
| Anti-Iba1 | FUJIFILM Wako Pure Chemical Corporation | 019-19741 |
| Donkey Anti-Rat AF488 | Thermofischer Scientific | A-21208 |
| Donkey Anti-Human Cy5 | Jackson ImmunoResearch | 709-175-149 |
| Donkey Anti-Human Cy3 | Jackson ImmunoResearch | 709-165-149 |
| Donkey Anti-Chicken Cy5 | Jackson ImmunoResearch | 703-175-155 |
| Donkey Anti-Rabbit Cy3 | Jackson ImmunoResearch | 711-165-152 |
| Donkey Anti-Rabbit AF488 | Thermofischer Scientific | A-21206 |
| Donkey Anti-Goat AF647 | Thermofischer Scientific | A-21447 |

Supplementary table 2: Antibodies used for flow cytometry

| Name | Clone | Company | Catalogue Number |
|---|---|---|---|
| BUV805 CD45 | 30-F11 | BD | 748370 |
| APC-efl780 CD45 | 30-F11 | eBioscience | 47-0451-82 |
| BUV395 Cd11b | M1/70 | BD | 563553 |
| BV421 CD64 | X54-5/7.1 | Biolegend | 139309 |
| BV711 CD64 | X54-5/7.1 | Biolegend | 139311 |
| Efl660 Lyve1 | ALY7 | eBioscience | 50-0443-82 |
| PE CD321 (Jam-A/F11r) | H202-106 | BD | 564908 |
| BV786 Ki67 | B56 | BD | 563756 |
| AF488 Cx3cr1 | SA011F11 | Biolegend | 149022 |
| BV605 Cx3cr1 | SA011F11 | Biolegend | 149027 |
| Efl450 CD3 | 17A2 | eBioscience | 48-0032-82 |
| APC-CY7 CD8a | 53-6.7 | Biolegend | 100714 |
| BV605 CD4 | RM4-5 | Biolegend | 100548 |
| PE-CY5 CD19 | eBio1D3 | eBioscience | 15-0193-82 |
| BV421 Cd49b | DX5 | BD | 563063 |
| APC TGFβ | 1D11 | R&D | IC1835A |

Supplementary table 3: Primer sequences

| **Gene** | **Forward primer** | **Reverse primer** |
|---|---|---|
| Rpl32 | AAGCGAAACTGGCGGAAAC | TAACCGATGTTGGGCATCAG |
| F11r | CTGTCCTGGTAACACTGATTCTC | GCAGTCCCTTTCTTTGTTCTTTC |
| Hexb | GACGACCAGTCTTTCCCTTATC | CCGGACATCGTTTGGTGTATAG |
| Cx3cr1 | TTCCCATCTGCTCAGGACCTC | AGACCGAACGTGAAGACGAG |
| Ccrl2 | GTCAACCCGCTGCTCTATTT | GCTGATAGCCTCCACTACTTTG |
| Tmem119 | GGATGCCTCACAGCTACAAA | TCCAACTCTGAGCCAATCATAAA |
| Olfml3 | CACCTTGTGGAGTACATGGAAC | CTACCTCCCTTTCAAGACGGT |
| Rgs1 | TTTTCTGCTAGCCCAAAGGA | TGTTTTCACGTCCATTCCAA |
| Uchl1 | GCCAGTGTCGGGTAGATGAC | TGGTTCACTGGAAAGGGCAT |
| Gfap | CCTGGAACAGCAAAACAAGGC | TTTCATCTTGGAGCTTCTGCCTC |

Supplementary video 1: Stitched time-lapse of ex-vivo live imaging of the muscularis externa of 14 day-old $Wnt1^{Cre/WT}$ $Rosa26^{tdT/WT}$ $Cx3cr1^{GFP/WT}$ mice. Scale bar is 20µm.

Supplementary video 2: Stitched time-lapse of ex-vivo live imaging of the muscularis externa of 56 day-old $Wnt1^{Cre/WT}$ $Rosa26^{tdT/WT}$ $Cx3cr1^{GFP/WT}$ mice. Scale bar is 20µm.
